# Experimental Test of Evolutionary Safety of a CRISPR-Cas9 Gene-Drive Element

**DOI:** 10.1101/2023.11.28.569142

**Authors:** Michael S. Overton, Sean E. Guy, Xingsen Chen, Alena Martsul, Krypton Carolino, Omar S. Akbari, Justin R. Meyer, Sergey Kryazhimskiy

## Abstract

CRISPR-Cas9 gene drives (CCGDs) are powerful tools for genetic control of wild populations, with applications from disease eradication to species conservation. However, Cas9 alone and in a complex with gRNA can cause double-stranded DNA breaks at off-target sites, which could increase the mutational load and lead to unintended loss-of-heterozygosity (LOH) events. These undesired effects raise potential concerns about the long-term evolutionary safety of CCGDs, but the magnitude of these effects is unknown. To measure how the presence of a CCGD or a Cas9 alone in the genome affects the rates of LOH events and de novo mutations, we carried out a mutation accumulation experiment in yeast *Saccharomyces cerevisiae*. We found no detectable effects on the genome-wide rates of mutations or LOH events. Our power calculations suggest that CCGD or Cas9 affect these rates by less than 30%, which is much less than natural variation for these traits in yeast. A more detailed examination shows that CCGD or Cas9 may alter the lengths and genomic distributions of LOH events, but the statistical support for these effects is weak. Thus, our results demonstrate that CCGDs impose at most a weak additional mutational burden in the yeast model. Although mutagenic effects of gene drives need to be further evaluated in other systems, our results add credence to the proposition that the evolutionary risks posed by well designed gene drives are likely acceptable.

## Introduction

The CRISPR-Cas9 based gene drives (CCGDs) are synthetic genetic elements that can rapidly spread in sexual populations through super-Mendelian inheritance^1^. The ability to encode various functions within CCGDs promises to give us an unprecedented degree of control over wild populations^2^. For example, by encoding a *Plasmodium*-disrupting peptide within a CCGD, it may be possible to reduce or even eliminate the spread of malaria^3–5^; a CCGD encoding drought resistance may allow us to prevent the extinction of populations vulnerable to climate change^2,6^. However, large-scale deployment of CCGDs in the wild faces several significant biological, as well as ethical, challenges^7–9^. Two major biological challenges are caused by evolution^7^. The first, short-term, problem is that certain mutations can arise in the CCGD element itself, abolishing its activity. Given that CCGD carriage often comes with a fitness cost, such loss-of-function mutations would be favored by selection and would result in gene drive-resistant populations^10–12^. This problem is widely recognized, and various engineering solutions have been proposed to mitigate it^13–15^. The second problem is that Cas9 has off-target effects^16–19^, which could increase the incidence of new mutations as well as loss-of-heterozygosity (LOH) events^20^ across the genome, potentially altering long-term evolutionary trajectories of treated populations. However, we know very little about the magnitude of these effects and their evolutionary significance^7,21^.

CCGDs vary in their design and complexity, but every CCGD contains a guide RNA (gRNA) that targets a particular genomic sequence (“the target allele”) and a Cas9 endonuclease^2,22^. In addition, CCGDs can also contain a genetic “payload” that produces a desired phenotype. The drive construct must be flanked by sequences homologous to those surrounding the target allele. When an engineered organism homozygous for the CCGD mates with a wildtype individual carrying the target allele, the fusion of gametes brings the drive and the target sequences together into the same cell. The Cas9/gRNA complex cuts the target allele, and the entire CCGD can be copied in its place by the homology directed repair (HDR) mechanism, thereby converting an initially heterozygous offspring into one homozygous for the CCGD allele. Through this process, CCGDs can achieve up to 98% inheritance, rapidly fixing the desired phenotype in the target population^4,5^.

The presence of a CCGD in the genome can change the rates and types of genome-wide mutations via several known as well as possibly other, as of yet unknown, mechanisms^16–19,23–25^. The best understood mechanism is template promiscuity whereby the Cas9/gRNA complex binds and cuts DNA sequences that are similar but not identical to the target^17,18,23,26^. These off-target dsDNA cuts are repaired by HDR or non-homologous end-joining (NHEJ), leading to LOH and indel mutations, respectively, at loci other than the target^16,23^. Off-target mutations may also be generated because the Cas9-gRNA complex bound to DNA might interfere with normal replisome progression. Importantly, such binding transiently occurs not only at the sequences similar to the target but also at random PAM sites^27^. Moreover, the Cas9 protein alone has a high, non-specific affinity for DNA^27^, which can potentially introduce additional mutations at non-PAM associated sites through the same interference mechanism. These additional mutations and LOH events could have unpredictable evolutionary consequences. For example, some LOH events could resolve hybrid incompatibilities and improve fitness^28–30^. Perhaps more likely, by reducing genetic diversity and exposing recessive deleterious alleles, LOH events could potentially lead to population decline, particularly in species that are already endangered^31,32^. However, the severity of these long-term evolutionary consequences depends on how strongly CCGDs affect the rates of mutations and LOH events, on the type of these events, and on their distribution along the genome. Thus, to quantify evolutionary risks associated with CCGDs, we need to measure how CCGDs affect the rates and the distributions of these events.

To address this problem, we designed a mutation accumulation (MA) experiment in yeast *Saccharomyces cerevisiae* with the aim to detect potential effects of a CCGD or Cas9 alone on the rates and genomic distributions of LOH events. While we also measure how these elements affect rates of point mutations and indels, our main focus is on LOH events because they result from the same molecular mechanism (HDR) that supports the intended activity of the CCGD. Importantly, MA experiments allow us to detect mutagenic effects caused by known as well as unknown molecular mechanisms. Finally, yeast is an established and powerful system for testing CCGDs^33^. Its major advantage is that we can carry out our experiments in a strain that is heterozygous at about 40,000 sites throughout the genome and maintain hundreds of independent MA lines for hundreds of generations^34–40^. These features maximize our statistical power to detect even relatively small changes in LOH rates caused by a CCGD or a Cas9 alone.

## Results

### Mutation accumulation experiment for detecting the effects of a CCGD on LOH rates

To measure the genome-wide rates of LOH events in the presence and absence of a CCGD element, we designed a mutation accumulation (MA) experiment using a hybrid-based scheme similar to that implemented in several recent studies^36–39,41^. The hybrid ancestors of our MA experiment (later referred to as “founders”) were generated by crossing the laboratory strain BY and the vineyard isolate RM (Table S1). The two parent strains differ from each other by approximately 40,000 SNPs and small indels spaced on average every 320 bp apart across the yeast genome^42^, such that the founders are heterozygous at these “marker” sites. An LOH event that occurs during the MA experiment can be detected if it converts one or more adjacent initially heterozygous markers to one of the parental homologs (Figure 1A). While we cannot detect very short LOH events, most of heterozygosity losses in yeast occur through large so-called terminal LOH events that typically extend over tens or hundreds of thousands of basepairs^34,35,38,39^. Therefore, the relatively high density of markers in our founders allows us to detect the vast majority of LOH events that contribute to heterozygosity losses in yeast.

**Figure 1.**
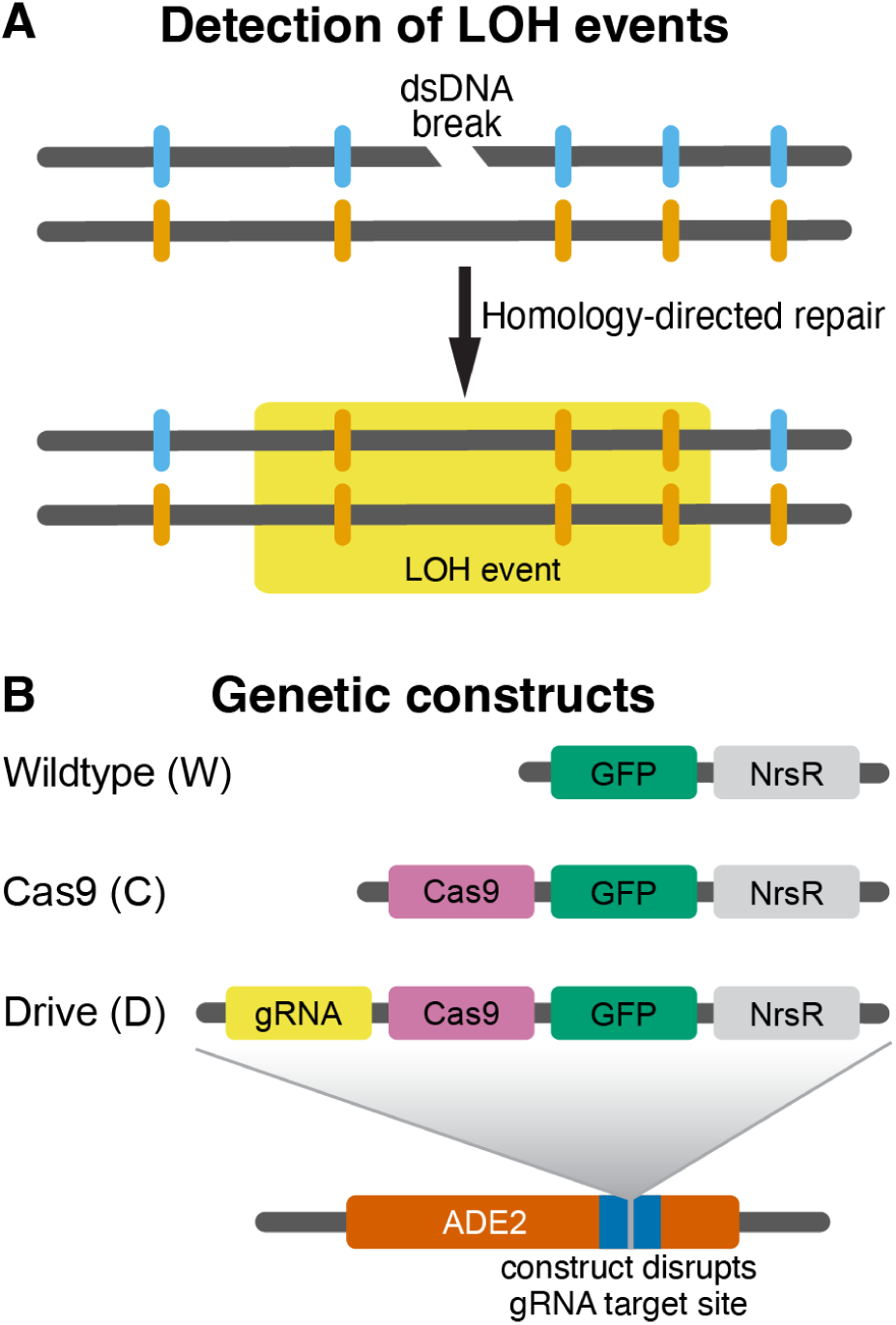
Schematic of experimental features. **A.** Detection of LOH events. Our founder strains are diploid hybrids heterozygous at about 40,000 marker sites across the genome (blue and orange notches). If a dsDNA break is repaired by homology directed repair, it causes loss of heterozygosity at nearby marker sites, which can be detected by whole-genome sequencing. **B.** Design of genetic constructs for the three founder strains. NrsR is the nourseothricin N-acetyl transferase gene which confers resistance to nourseothricin. All constructs are integrated into the ADE2 locus at Chr XV and disrupt the gRNA binding site within this gene.

We constructed three types of founder hybrids by integrating different constructs into the ADE2 locus (Figure 1B, Table S1; see Section “<u>Strain construction</u>” in Materials and Methods). The “drive” founders (or “D” for short) are homozygous for the constitutively expressed CCGD with the gRNA targeting ADE2 (Figure 1B). Since the ADE2 gene is disrupted by the integrated CCGD cassette, the target sequence is absent from the genome of D strains, such that any cut performed by the Cas9/gRNA complex would be off-target. This type of strain was designed to simulate a gene drive that had successfully fixed in a population, and is now permanently integrated into the species’ genome. The “Cas9” founders (or “C” for short) are homozygous for the constitutively expressed Cas9 gene. We designed this type of strain to look for possible mutagenic effects of a “naked” Cas9 protein, i.e., one that is not associated with any gRNAs, something that can occur for example if the Cas9 and gRNA expression levels are not perfectly matched^18^. Finally, the wildtype control founders (or “W” for short) are homozygous for the integration cassette without any gene-drive components.

Our a priori expectation was that the main potential side-effect of the presence of a CCGD element in the genome would be an HDR-driven increase in the rate of LOH events at off-target sites. Previous studies reported up to about 50-fold variation in LOH event rates among yeast hybrids^36,38–40^. Since we anticipated that the CCGD-associated effects would likely be smaller, we sought to design a sufficiently powerful MA experiment to detect less than two-fold changes in LOH rates. Modeling the accumulation of LOH events in each MA line as a Poisson process we found that, if each wildtype MA line accumulates on average 13 LOH events, the probability of detecting a 15% increase in the LOH rate with 95 MA lines per founder at P-value of 0.01 would be 85.7% (Figure 2A; see Section “<u>Analysis of</u> <u>statistical power</u>” in the Materials and Methods). Given that the previously measured rate of LOH events averages 2.3×10^−2^ (range from 3.6×10^−3^ to 4.7×10^−2^) per genome per generation^36–40,43,44^, we expected to achieve this statistical power by propagating our MA lines for 750 cell divisions.

**Figure 2.**
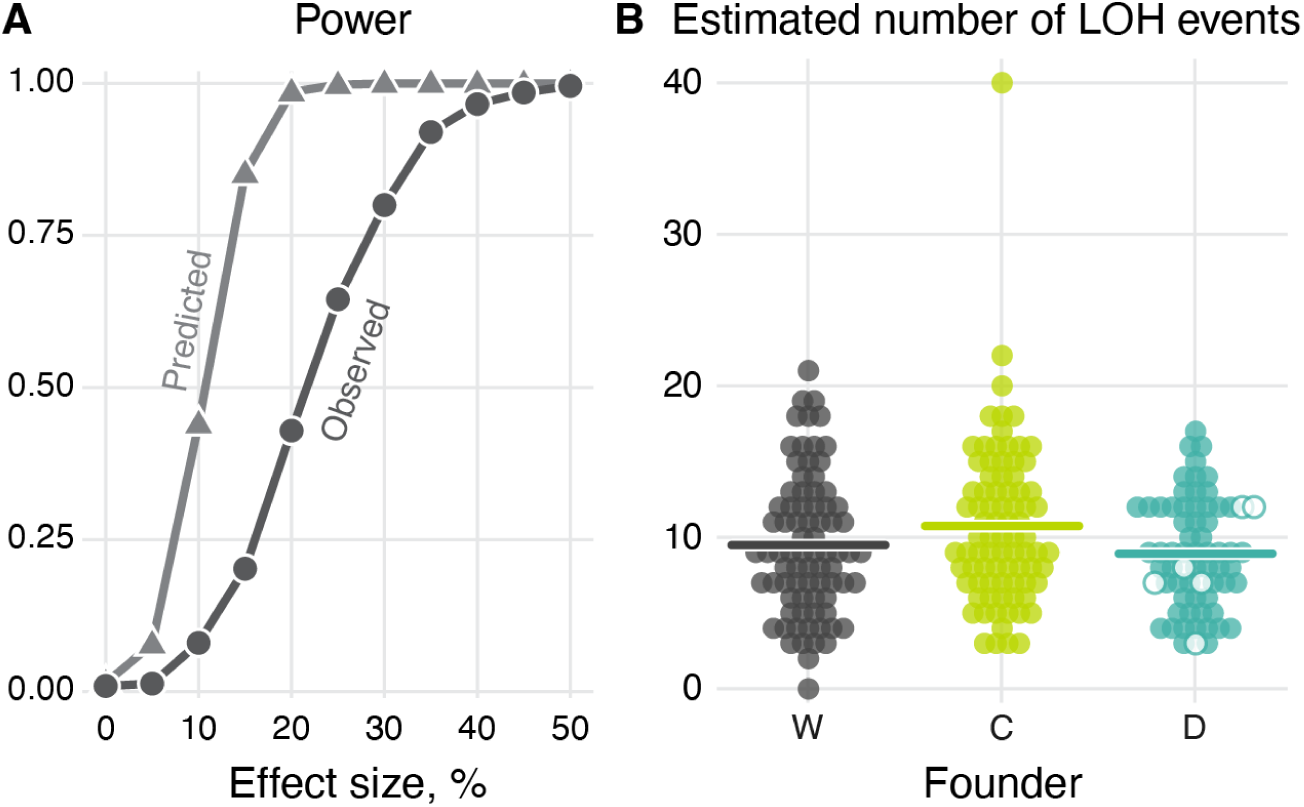
Statistical power to detect differences in the LOH rates and the estimated number of LOH events in our mutation accumulation experiment. **A.** The theoretically predicted and the observed power to detect a certain percentage increase in the LOH rate at *P*-value 0.01 (see text for details). **B.** The number of the LOH events per end-point clone of each founder after correcting for undetected events. Hollow circles indicate clones tested for CCGD activity (see <u>Figure S1A</u> and Section “<u>Testing of gene-drive activity</u>” in the Materials and Methods). Horizontal lines indicate the mean number of LOH events per clone.

Guided by these a priori estimates, we isolated 8, 8 and 7 founder clones of types D, C and W, respectively and established between 11 and 14 MA lines per clone, which resulted in 95 MA lines per D, C and W type each, for a total of 285 MA lines. We propagated these lines using a standard MA protocol^45^ (see Section “<u>Mutation</u> <u>accumulation experiment</u>” in the Materials and Methods). Briefly, for each MA line, we picked a random colony from a previous growth cycle, streaked it to single cells and let the resulting colonies grow for 48 hours. Assuming that each growth cycle corresponds to 20 cell divisions, we predicted that a typical MA line would accumulate 13 LOH events after 38 cycles. To account for variation among LOH-rate estimates across studies, we propagated our MA lines for 43 cycles. After the experiment was completed, we counted the number of cells per colony after 48 hours of growth and estimated that our MA lines in fact underwent approximately 800 cell divisions (see Section “<u>Mutation accumulation experiment</u>” in Materials and Methods). We then sequenced the full genomes of all founder and end-point clones to a median depth of 34.2×. After excluding clones that did not pass our quality-control checks (see Section “<u>Analysis of sequencing data</u>” in Materials and Methods), we retained 79, 83, and 67 end-point clones for subsequent analyses from W, C, and D lines respectively.

Before estimating LOH rates from these data, we used two approaches to verify that the CCGD element remained active during the experiment. First, we picked six end-point D clones and experimentally confirmed that their CCGD element was in fact active (see Section “<u>Testing gene-drive activity</u>” in Materials and Methods and <u>Figure S1A</u>). Second, we used our whole-genome sequencing data to identify any de novo mutations that occurred within our constructs and could have disrupted the CCGD function. We found a total of six end-point clones (1 W, 3 C, and 2 D clones) with such mutations, with each clone carrying either a single heterozygous SNP or indel within the construct (<u>Figure S1B</u>). Only two of these mutations occurred in the drive components, a non-synonymous SNP (A299T) in the Cas9 gene of a C clone and an indel at the 3’ end of the gRNA promoter in a D clone. The number of observed construct mutations is roughly consistent with our expectation (see Section “<u>Estimating the expected number of mutations in the construct</u>” in Materials Methods). We conclude that even if all of these mutations deactivated the gene-drive components, they would only minimally decrease our power to detect differences in LOH rates between founders.

In summary, we designed and carried out an MA experiment with a reasonable statistical power to detect possible effects of a gene drive and/or naked Cas9 on the rates of LOH events. All available evidence strongly suggests that the gene-drive components in our D and C MA lines remained active during this experiment.

### Upper bound on the increase in the genome-wide LOH-rate imposed by CCGD or Cas9

We next used a novel dual-reference pipeline that we developed^46^ to genotype our end-point and founder clones and identify LOH events. We discovered a total of 1,645 LOH events across all of our 229 our MA lines (a median of 7 per end-point clone), supported by a total of 279,788 converted markers (Table S2). We detected on average 648/83 = 7.8 and 444/67 = 6.6 LOH events per C and D line, respectively (<u>Figure S2</u>, Table 1), which is statistically indistinguishable from 553/79 = 7.0 events detected in a typical W line (*P* = 0.131 and P = 0.419 for C and D lines respectively; permutation test).

**Table 1.** Key mutation counts and rates among end-point clones from the three strains.

|  | Loss of Heterozygosity |  |  |  |  |  |  | Point mutations |  | Indels |  |
| --- | --- | --- | --- | --- | --- | --- | --- | --- | --- | --- | --- |
|  | End-point clones | Detected events | Estimated events | Event rate<sup>1</sup><br>×10<sup>-2</sup> | Conv. rate<sup>2</sup><br>×10<sup>-5</sup> | Estimated iLOH | Estimated tLOH | Total | Rate<sup>2</sup>, ×10<sup>-11</sup> | Total | Rate<sup>2</sup> ×10<sup>-11</sup> |
| W | 79 | 553 | 785 | 1.24 ± 0.13 | 7.61 ± 1.36 | 625 | 160 | 70 | 5.35 ± 1.45 | 48 | 3.67 ± 0.99 |
| C | 83 | 648 | 936 | 1.40 ± 0.16 | 6.11 ± 1.03 | 784 | 153 | 77 | 5.61 ± 1.53 | 43 | 3.13 ± 1.16 |
| D | 67 | 444 | 627 | 1.16 ± 0.12 | 6.73 ± 1.26 | 490 | 137 | 53 | 4.78 ± 1.26 | 45 | 4.06 ± 1.17 |

<sup>1</sup>per genome per generation ± 95% bootstrap CI
<sup>2</sup>per base-pair per generation ± 95% bootstrap CI

After correcting for undetected events (see Section “<u>Analysis of sequencing data</u>” in the Materials and Methods and Ref. ^46^), we estimate that a total of 2,348 LOH events with length exceeding 17 bp likely occurred across our 229 MA lines. We estimate that a typical C and D line accumulated 936/83 = 11.2 and 627/67 = 9.35 of LOH events, respectively (Figure 2B and Table 1), statistically indistinguishable from 785/79 = 9.94 events accumulated by a typical W line (*P* = 0.131 and P = 0.419 for C and D lines respectively; permutation test).

On average, 5.5% of the genome per clone lost heterozygosity during our MA experiment. This fraction was 4.9% and 5.4% for a typical C and D clone, respectively, statistically indistinguishable from 6.1% for a typical W clone (*P* = 0.205 and *P* = 0.652, permutation test; Table 1, <u>Figure S3</u>).

Given the absence of a detectable effect of the CCGD or Cas9 on the overall LOH event rates, we sought to estimate the upper bounds on these effects given the power of our experiment. Since the estimated numbers of LOH events that occurred during our experiment are somewhat lower than predicted theoretically and have greater variance than a Poisson distribution (<u>Figure S4</u>), the statistical power of our experiment is slightly reduced. When we re-estimate the power of our experiment, the chance of detecting a 15% LOH-rate increase at the P-value 0.01 in our data is in fact about 20% instead of the anticipated 85.2% (Figure 2A; see Section “<u>Analysis of statistical power</u>” in Materials and Methods for details). Nevertheless, our experiment gives us power close to 100% to detect the effect sizes above 45% and substantial power to detect the effect size above 30%. Therefore, our results indicate that the presence of a CCGD or the Cas9 protein alone increases the LOH rates in yeast (if at all) likely by less than 30% and almost certainly by less than 45%.

### Effects of CCGD and Cas9 on LOH characteristics and their genomic distribution

#### Interstitial versus terminal LOH events

Double-stranded DNA breaks can be repaired by multiple HDR mechanisms^47–50^, resulting in the so-called “interstitial” and “terminal” LOH events (iLOH and tLOH). In particular, tLOH events are associated with replication stalling during S-phase and may occur due to replisome collisions with protein-DNA complexes^51,52^. Since the DNA-binding activity of Cas9, alone or in complex with a gRNA, could interfere with the DNA processing during HDR or produce replication fork stalling, it is possible that the presence of these elements could change the rates of iLOH and tLOH events differently even if the total LOH rate changes are undetectable. To test for this possibility, we classified all 2,348 inferred LOH events into 1,898 (80.8%) interstitial and 450 (19.2%) terminal events (see Section “<u>Analysis of sequencing data</u>” in Materials and Methods). We found that the C and D strains accumulated on average 1.84 and 2.04 tLOH events, respectively, which is statistically indistinguishable from 2.03 tLOH events accumulated by the W strain (*P* = 0.389 and *P* = 0.950, respectively, permutation test; Table 1). The C and D clones accumulated on average 9.44 and 7.31 iLOH events, which again were statistically indistinguishable from the 7.91 events found in a typical W clone (P = 0.056 and P = 0.373, respectively).

#### LOH length distributions

Interactions between Cas9, with or without a gRNA, and genomic DNA could also disrupt key repair processes such as strand resection, strand invasion, or polymerase progression, which could potentially lead to changes in the length distributions of ensuing LOH events. We find that the median length of an iLOH event is 430 and 752 bp in C and D backgrounds (<u>Figure S5</u>), which is statistically indistinguishable from 471 bp in the W background (*P* = 0.544 and P = 0.207; permutation test; see Section “<u>Comparison of LOH rates across strains</u>” in Materials and Methods). However, tLOH events are shorter in both C (median = 131 kb, *P* = 0.034) and D (median = 142 kb, *P* = 0.133) strains than in the W control (median = 213 kb).

#### Distribution of LOH events across the genome

Given that the CCGD off-target activity is sequence specific, it is possible that CCGD or the naked Cas9 protein strongly affect LOH rates only in certain regions of the genome and that such local effects are missed when the data are pooled across the whole genome. Thus, we next looked for possible effects of these elements at the chromosome-arm and 50-kb scales, which trade-off statistical power with resolution. We found no difference in the distributions of tLOH events across strains at either scale (<u>Figure</u> <u>S6</u>). At the same time, while the distributions of iLOH events are indistinguishable between strains at the 50-kb scale (P > 0.074, Wasserstein test after Benjamini-Hochberg correction; Figure 3B), they are somewhat different at the chromosome-arm scale (*P* = 0.047 and 0.036 for C and D strains compared to W; χ^2^ test; Figure 3A). However, we could not localize these differences to any specific chromosome arms (P > 0.192, permutation test after Benjamini-Hochberg correction), nor did we find any detectable effect of this difference on the distribution of LOH conversions across the genome (<u>Figure S7</u>).

**Figure 3.**
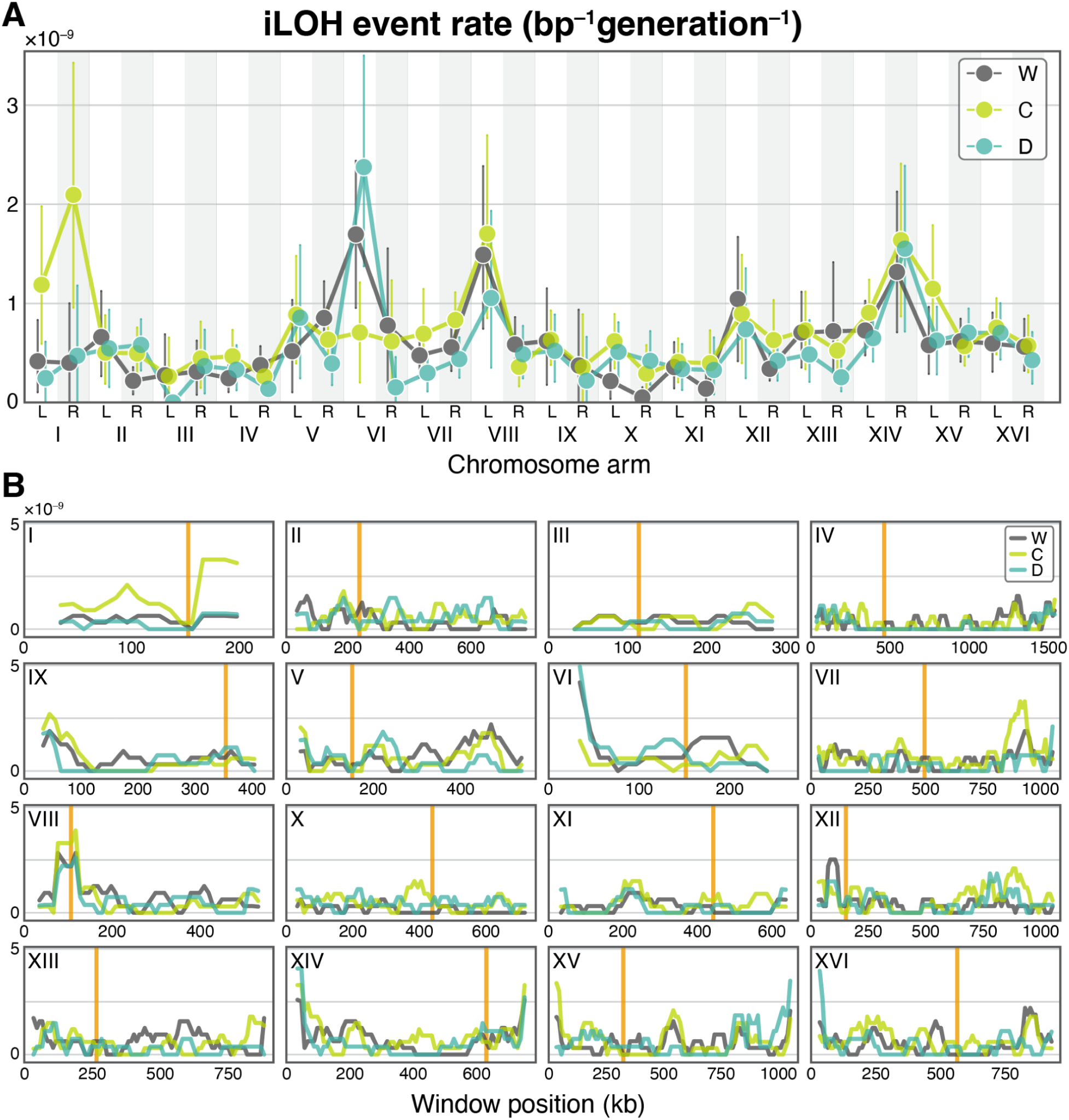
Distribution of iLOH event rates across the genome. **A.** iLOH event rates across chromosome arms. Vertical bars indicate the 95% bootstrap confidence intervals. **B.** iLOH event rates across 50 kb sliding windows on each chromosome with a 10 kb step size. Orange vertical lines indicate centromeres.

We also tested whether LOH event rate variation is associated with certain nucleotide sequences. To this end, we scanned the BY and RM genomes for PAM sites and scored the similarity of the 5’-adjacent sequences to our gRNA (see Section “<u>Comparison of LOH rates across strains</u>” in Materials Methods). We found no evidence that the LOH event rates were elevated in C or D backgrounds in those 50 kb windows that harbored more motifs similar to our gRNA (<u>Figure S8A</u>) or those where densities of PAM sites were higher (P > 0.558, linear regression; <u>Figure</u> <u>S8B,C</u>).

Taken together, these results suggest that, to the first approximation, the LOH characteristics and their distribution along the genome are largely unaffected by the presence of a CCGD or naked Cas9. However, the full picture is more nuanced. We carried out several statistical comparisons and found three with P-values slightly below 0.05, with the three being the effect of Cas9 on the lengths of tLOH events and the effects of CCGD and Cas9 on the genome distribution of iLOH events. Given that these results are only marginally significant, we cannot rule out the possibility that they represent Type I errors. But the possibility that CCGD and Cas9 elements slightly change some LOH characteristics and their distribution also remains.

### Upper bound on the increases in the genome-wide mutation rates imposed by CCGD or Cas9

In addition to affecting LOH rates, the presence of the CCGD or Cas9 protein could also increase the rate of new mutations. To test this, we identified new mutations in our end-point clones (Table S3; see Section “<u>Analysis of sequencing data</u>” in Materials and Methods). We found a total of 200 single-nucleotide mutations (SNMs) and 136 small indels (< 50 bp) across all end-point clones. A typical C and D end-point clone carries 120/83 = 1.45 and 98/67 = 1.46 SNMs or indels, which is statistically indistinguishable from 118/79 = 1.49 mutations found in a typical W clone (Figure 4, Table 1; P = 0.843 and P = 0.629 for C and D clones respectively; permutation test). Estimating the statistical power of our experiment to detect differences in mutation rates, we found that a detection of a 50% increase in mutation rate would have been almost certain, and a 30% increase would have been detected with probability 80% (Figure 4A). Thus, we conclude that presence of an active CCGD or naked Cas9 increases point mutation rates in yeast (if at all) likely by less than 30% and almost certainly by less than 50%.

**Figure 4.**
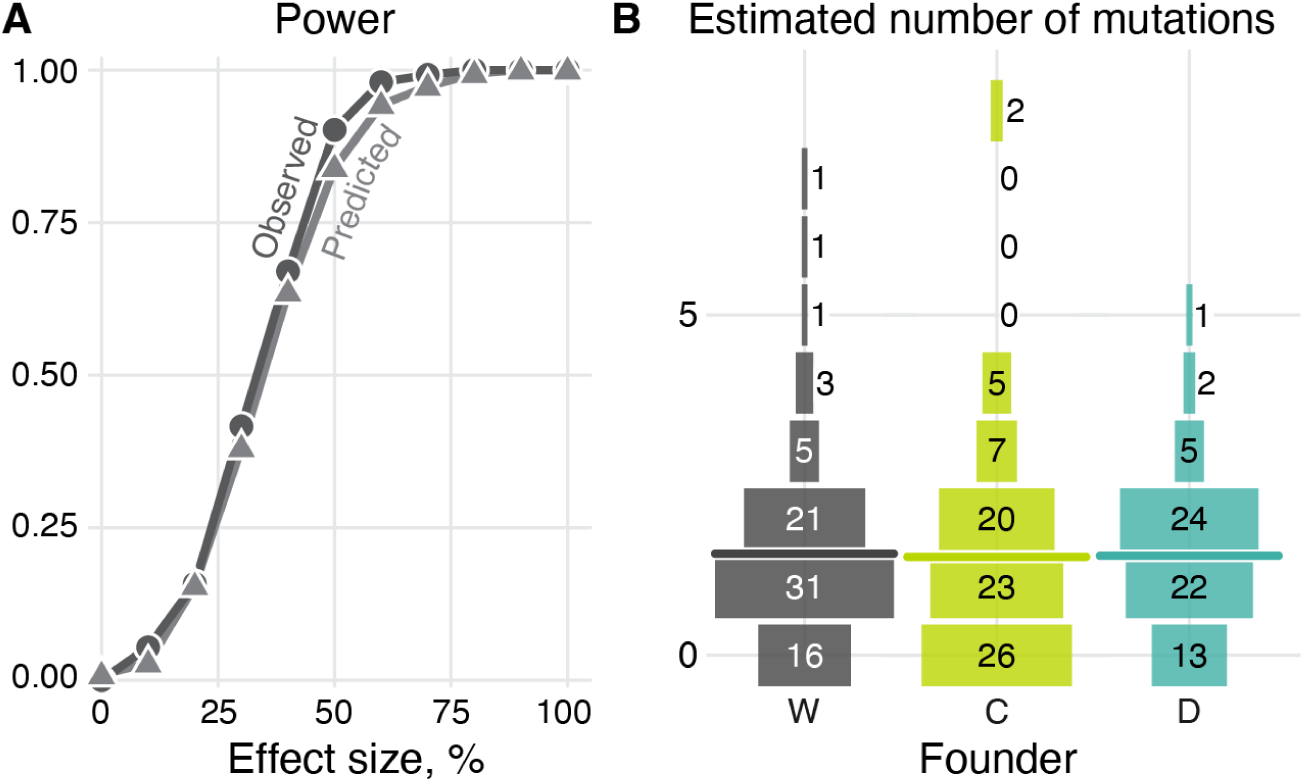
Statistical power to detect differences in the mutation rates and the estimated number of mutations in our mutation accumulation experiment. **A.** Same as Figure 2A but for mutations (both single nucleotide variants and small indels are included). **B.** Distribution of mutations per clone of each strain. Horizontal lines indicate the mean.

## Discussion

In this study, we investigated how the presence of a CRISPR-Cas9 gene drive (CCGD) element or the Cas9 gene without a dedicated guide RNA in the genome affects the rates of LOH events and new mutations in yeast. We found no evidence that the overall LOH or mutation rates were elevated in strains carrying the CCGD or the Cas9 gene. Given the large size of our study, the absence of statistically significant effects on the overall LOH rate strongly suggests that these effects are small. Specifically, we estimate that the presence of a CCGD or the Cas9 protein alone increases the LOH and mutation rates likely by no more than 30%. By comparison, previous studies found up to a 50-fold difference in the LOH event rates among yeast hybrid strains^40,46^. Thus, the possibility of a 30% increase in the LOH event rate due to CCGD carriage is quite modest compared to that produced by natural variation. Similarly, an increase in the point mutation rate of 30% is much smaller than the five-fold variation in mutation rates among yeast strains^39^ and the six-fold variation observed across the genome of a single yeast strain^53^.

While we find no detectable effects of CCGD or Cas9 on the genome-wide LOH rates, the presence of Cas9 apparently makes tLOH events shorter. Given that this result is only marginally significant (*P* = 0.034), we cannot rule out a Type I error. But this shift could also point to a real biological effect whereby the presence of Cas9 alters the probabilities with which the cell engages different DNA repair mechanisms, such that, for example, dsDNA breaks that occur closer to centromeres are more likely to result in iLOH events in C strains than in W strains. Consistent with this hypothesis, C strains appear to accumulate iLOH events about 20% faster than controls (although this difference is not statistically significant).

We found a similarly weak statistical support for the effects of CCGD and Cas9 on the genome distribution of iLOH events. These could also be Type I errors, or they could indicate that Cas9, alone or in a complex with gRNA, alters where in the genome dsDNA breaks occur or which repair mechanisms are engaged. Given that the Cas9:gRNA complex is known to produce off-target dsDNA cuts at sequences that display sufficient complementarity to the gRNA^16–19,25,26^, we anticipated that a CCGD element would increase LOH event rates at loci that are most similar to our gRNA. However, we found no sequence-specific local effects, although our power to detect them is quite low. Consistent with our observations, another study found a similar lack of clear off-target effects in mosquitos carrying well-designed gene-drives^21^. Importantly, our gRNA was designed so that all non-target PAM-adjacent genomic sequences contain a minimum of 7 mismatches against our gRNA^33^. Previous studies have shown that cutting activity at such sites is negligible^18,26,33^. Thus, our results are consistent with the expectation that current off-target prediction algorithms likely capture all the strongest off-target sites, although a possibility that they miss weaker (and probably more broadly distributed) sites remains.

We propagated our yeast MA lines asexually, and the CCGD and the Cas9 gene were expressed constitutively. However, CCGDs are often designed to be expressed only transiently during meiosis. Given that the mechanisms of the activities of various DNA repair mechanisms differ between meiosis and mitosis^48^, it is possible that the mutagenic effects of such CCGDs may be different from those reported here. However, we expect that the overall mutagenic effect would be even weaker than in our system simply because the genome in such systems would be less exposed to Cas9.

Overall, our main finding that a CCGD with a well-designed gRNA or the Cas9 alone induce at most weak mutagenic effects suggests that the long-term evolutionary risks associated with CCGD carriage are probably acceptable. However, one should keep in mind that our results were obtained in a unicellular model organism under lab conditions. It is a priori unclear how they will translate to multicellular organisms in natural environments. In addition, to measure the effects of CCGDs or Cas9 on mutation rates, we purposefully eliminated all but the strongest selection pressures. However, even weak selection pressures imposed on the organism by the presence of a CCGD element may be sufficient to change its long-term evolutionary trajectory. Such selection pressures may be very hard to measure and even harder to predict a priori, since they may be highly variable depending on the payload gene, the target species and the environment. Thus, further investigations will be needed to further assess evolutionary risks of CCGDs. These considerations notwithstanding, our results reaffirm that CCGDs are a viable option for population control.

## Methods

### Experimental methods

#### Strain construction

All strains are listed in Table S1. We generated the W, C and D strains by mating two haploid yeast strains. One parent strain is derived from the laboratory *S. cerevisiae* strain BY4741 (BY; MATa his3Δ1 leu2Δ0 met15Δ0 ura3Δ0) and the other parent is derived from the vineyard strain YAN501 (RM; MATα his3Δ1 leu2Δ0 ura3Δ0 HO::KanMX)^54^. These two parent strains differ by ∼40,000 SNPs and small indels^42^. We used standard transformation techniques to replace the his3Δ1 allele in BY parent with a functional version to facilitate diploid selection^55^. We amplified the NatMX-GFP, NatMX-GFP-Cas9 and NatMX-GFP-Cas9-gRNA constructs from a previously constructed gene drive cassette, p414-Cas9 and p426-gRNA[ADE2]^33,56^ and integrated each of them into each of the parent strains at the ADE2 locus (Chr XV:564476-566191) using standard yeast transformation methods and selection on YPD+nourseothricin (nat) plates. The gRNA sequence used here targets the 5’ end of the ADE2 gene, and the disruption of ADE2 produces an easily observed red colony phenotype^33^. Single colonies of BY and RM haploid parents containing each of the integrated constructs were mated, diploids were selected on G418+his– plates, and correct diploid progeny were confirmed by PCR targeting the Ade2 region. Diploid founders were each derived from independently transformed and mated BY and RM parents to reduce the chance of multiple lines being established from a mutated founder. To test the activity of our CCGD, we also constructed two Mat**a** “tester” haploids derived from strain MJM64^57^ by knocking in one of two functional copies of ADE2. MJM64 was transformed with the BY4742 copy of ADE2 to generate the intact target strain (IT), yKC57, and with the YAC2485^33^ copy of ADE2 sequence to generate the no target (NT) strain yKC29. The YAC2485 ADE2 sequence is engineered with 12 mismatches to abolish gRNA targeting. In both tester strains, URA3 is driven by the haploid-specific promoter STEP5pr which makes them 5-FOA sensitive as haploids but not as diploids. They also carry a KanMX cassette, which makes their progeny G418 resistant.

#### Mutation accumulation experiment

For each of the three constructs, we used eight founders to establish 95 independent mutation accumulation lines, for a total of 285 MA lines. Each of these lines was independently propagated for 43 cycles of streaking to single colony isolation on YPD agar plates and incubating at 30°C for 48 h, for a total average of 800 generations. The colony closest to a pre-marked location on the plate was used to continue the next cycle to prevent any potential artificial selection effects. Founder and end-point clones were preserved by culturing single colonies in YPD liquid media to mid-log growth and freezing at –80°C in 15% glycerol. To estimate the number of elapsed generations, we grew one founder and 10 end-point clones from each of the W, C and D backgrounds under identical conditions to the MA experiment in four biological replicates. For each replicate, we picked a colony, resuspended it in PBS, estimated the cell count using the Coulter Counter, and calculated the number of generations within a cycle as log_2_(colony size).

#### Testing of gene-drive activity

To test whether the gene drive elements remained active by the end of the MA experiment, we sampled six end-point D clones, three exhibiting low LOH accumulation and three chosen randomly. We sporulated and dissected tetrads for each diploid clone and selected Matα haploids that were -ura and -KanMX, making them 5-Fluoroortic acid (5FOA) resistant and G418 sensitive. All Mat types of all haploids were confirmed with creep assays against strains MJM36 (Matα) and MJM64 (Mat**a**)^57^. Since 5-FOA and G418 require different media pH, we first selected clones on Complete Synthetic Media (CSM) +5FOA pH 4.5, then on CSM+G418 pH 7.0. For each original D clone, we picked one Matα haploid and mated it with the two Mat**a** “tester” haploids. After mating, we select for diploids using 5-FOA and G418 selection, as detailed above. After 48 h of growth on CSM+G418, colonies were photographed. The presence of red colonies indicates successful ADE2 disruption by an active CCGD^33,56^.

#### Library preparation and sequencing

Genomic DNA was extracted and purified from each founder and end-point clone using a modified ethanol precipitation protocol. Briefly, overnight cultures were pelleted and resuspended in 6% SENT (6% SDS, 10mM EDTA, 30mM Tris, pH8) and incubated at 65°C for 15 min. Cooled lysates were combined with RNAse A and incubated at 37°C for 60 min. A half volume of 3M NaOAc was added and the mixture was centrifuged to pellet proteins and cellular components. The supernatant was washed first in isopropanol and then ice-cold ethanol. DNA pellets were air dried in inverted tubes and resuspended in nuclease-free water. We prepared whole-genome sequencing libraries using the Illumina Nextera system with a modified protocol^58^. We pooled 100bp paired-end libraries and sequenced them on an Illumina Hi-Seq 4000 platform to a median read depth of 34.2x.

#### Gene drive safety

All work involving gene drive constructs was performed according to protocols approved by the Institutional Biosafety Committee at UCSD.

### Data analysis

#### Analysis of statistical power

##### Theoretically predicted power

To determine the statistical power of our MA experiment to detect differences in the rates of LOH events between strains, we modeled their accumulation as a Poisson process with rate λ = 1.76×10^−2^ per genome per generation^36,39^. Then, assuming that our experiment would take about *T* = 750 generations, the number of LOH events in each end-point clone is a Poisson random number with expectation µ = λT = 13.2. Given that we had the resources to maintain the maximum of 95 MA lines per founder, we simulated our experiment as follows. We first formed the “baseline” sample by drawing 95 random numbers from the Poisson distribution with mean µ = 13.2. We then generated the “alternate” sample of end-point clones whose event counts were drawn from the Poisson distribution with mean µ’ = µ(1 + 𝑓_𝑒_) where *f_e_* > 0 is the effect size. We then compared the differences in the means of these two samples using a permutation test with 1,000 independent permutations, and called the difference significant if the P-value was at or below 0.01. For each effect size *f_e_* between 5% and 50% with the step-size of 5%, we repeated this procedure 1,000 times and estimated the power of our experiment as the fraction of simulations that yielded a significant test.

##### Observed power estimation

We estimate the actual statistical power of our experiment as follows. We first pool all 229 of our end-point clones that carry a total of *n* = 2,248 LOH events to form the “baseline” sampling population. Then, at each realization of our simulation for a given effect size *f_e_*, we form the “alternate” population by randomly distributing *n*(1 + *f_e_*) additional LOH events (rounded to the nearest integer) across all 229 clones. We then randomly sample 79 clones from the baseline population and 67 clones from the alternate population and compare their sample means using the same permutation test as above. We carry out 1000 realizations at each effect size *f_e_* between 5% and 50% with the step-size of 5%, and obtain an estimate of power as above. The observed statistical power for point mutations was estimated analogously.

#### Estimating the expected number of mutations in the construct

Since our MA lines accumulated mutations for approximately 800 generations, it is possible that some mutations occurred in the construct and some of them may have deactivated the gene drive element. SNP and indels occur on average at rates of 1.67×10^-10^ and 7.5×10^-12^ per base pair per generation, respectively^39,59,60^. Given that our constructs D, C and W constructs are 7999 bp, 7601 bp and 2770 bp long, respectively, a typical end-point clone from each respective strain is expected to have 2.23×10^−3^, 2.12×10^−3^, and 7.73×10^−4^ mutations in the construct. Thus we expect a total of 0.15, 0.18 and 0.061 D, C and W end-point clones across the entire experiment to carry a mutation in the construct. Given that the length of the gRNA and Cas9 genes are 388 and 4890 bp (including promoter and terminator), respectively, we can conservatively estimate the fractions of gene-drive/Cas9 deactivating mutations as 66.0% and 64.3% in the D and C clones respectively. Thus, we expect 0.090 and 0.11 of the D and C clones to have a gene-drive or Cas9 deactivating mutation, respectively.

These estimates are based on the reasonable assumption that all mutations within the constructs are effectively neutral in our MA experiment, i.e., they do not provide a selective advantage or disadvantage substantial enough to overcome the strong genetic drift imposed by periodic single-cell bottlenecks. A more important source of uncertainty in these estimates arise from the fact that they are based on genome-wide average mutation rate estimates, but the actual mutation rates can vary up to 6-fold across locations in the genome^53^. Thus, we might reasonably expect anywhere between 0.065 and 2.34 out of 229 sequenced clones to carry a mutation in the construct. Similarly, we expect between 0.015 and 0.54 of the D clones to have a gene-drive-deactivating mutation and between 0.018 and 0.66 of the C clones to have a Cas9-deactivating mutation.

#### Analysis of sequencing data

Sequencing library preparation failed for five founders (H_F00, H_H00, N_F00, N_H00, F_D00). In order to use their descendant MA lines for LOH and mutation analyses, we imputed the ancestral genotypes from descendant end-point clones, as described below. Inclusion of these lines did not change our results. In addition, we found that the ancestors of two D MA lines (2 founders and 27 end-point clones) were apparently triploid as detailed in Ref. ^46^, compromising our ability to detect new LOH events, and were excluded. We excluded 19 additional clones from further analysis due to low coverage or traces of cross-contamination during library preparation.

##### Genotyping

We genotyped our founder and end-point clones at the initially heterozygous marker sites using our newly developed reference-symmetric genotyping pipeline^46^. Briefly, we align trimmed reads to both the BY and RM parental reference genomes. We exclude 1,774,809 of repeat regions identified by RepeatMasker^61^ and 7.5 kb at chromosome ends^54^, as all of these regions are difficult to map unambiguously. This reduces the length of the genome that we monitor for the presence of LOH events to an effective length of 10.3 Mb. To obtain genotype calls, we first use the GATK HaplotypeCaller^62–65^ with the BY and the RM reference genomes separately, reconcile the emitted calls at each marker site, and then remove dubious heterozygous calls.

For each of the four founders whose genomes were not sequenced (see above), we impute the genotype at a marker site as heterozygous (homozygous for a given allele) if at least six of its descendant end-point clones and all other founders genotyped at that site are called heterozygous (homozygous for the same allele).

##### Detection and exclusion of aneuploidies

We detect aneuploidy events as described in Ref. ^46^. We found no aneuploid events in the founders, but detected a total of 21 aneuploidies (12 chromosome losses and 9 chromosome gains) in 17 end-point clones, with 5, 8, and 6 events affecting the W, C and D strains, respectively. All aneuploid chromosomes were excluded from LOH analyses.

##### Identification of LOH events

To identify LOH events, we first find the so-called “LOH tracts” in an end-point clone as sequences of adjacent marker sites that are called homozygous for the same allele. A boundary of an LOH tract is estimated as the midpoint between its first converted marker and the adjacent unconverted marker (or a marker converted to a different homolog). We merge multiple LOH tracts into a single LOH event if the LOH-tract boundaries are separated by less than 10 kb^36^. We classify those LOH events that contain the last marker on a chromosome arm as terminal (tLOH); other LOH events are classified as interstitial (iLOH). The set of detected LOH events is listed in Table S2.

##### Corrections for undetected LOH events

After detecting LOH events and identifying their lengths, we apply three corrections for undetected iLOH and tLOH events: (i) correction for inter-marker iLOH events; (ii) correction for iLOH events overlapping with tLOH events; (ii) correction for overlapping tLOH events. These corrections are described in detail in Ref. ^46^. We compare both detected (i.e., uncorrected) and corrected LOH counts across strains, as indicated.

##### Estimation of causal dsDNA breakpoint positions

We estimate the positions of dsDNA breaks that likely led to the formation of an LOH event as follows. For iLOH events and tLOH events under 20 kb, we locate the breakpoint as the midpoint between its boundaries^47^. For tLOH events over 20 kb, we locate the breakpoint inside the tLOH event, 10 kb from its centromere-proximal boundary^36^. However, we note that this estimate is not very precise because the degree of strand resection—and therefore the length of the LOH region extending from the breakpoint—differs between the two dsDNA repair mechanisms (strand crossover during double-strand break repair or break induced-replication repair) that can produce tLOH events^50,66^.

##### Detection of de novo mutations

To detect de novo single-nucleotide mutations (SNM) and indels we start with the set of genotypes emitted by our genotyping pipeline and exclude all sites that differ between the two parental reference sequences or sites that we identified as ancestrally heterozygous. The remaining variants form the set of putative mutations. This set contains many sites with multiple identical base substitutions in individual clones. Given that the mutation rate in yeast is about 1.67×10^−10^ (Ref. ^59^), the expected number of mutations that occur at a given site during the entire MA experiment in any of the clones is µ = 1.67×10^−10^ × 800 × 229 = 3.06×10^−5^. Therefore, the probability that two or more mutations occur at a given site is P_2+_ = 1 – *e*^−µ^(1 + µ) ≈ µ^2^/2 = 4.68×10^−10^, so that we would expect 2*L*P_2+_ = 2 × 1.2×10^7^ × 4.68×10^−10^ = 1.12×10^−2^ sites across the whole genome to be mutated twice or more in the entire experiment, where *L* = 1.2×10^7^ is the haploid genome length of *S. cerevisiae*. Since indel rates are generally much lower than those of SNMs, about 5.0×10^-12^ per base pair per generation^59^, it is extremely unlikely that any duplicate SNMs or indels have occurred in our experiment. Therefore, we identify only unique variants (those that are present only in a single end-point clone) as new mutations. We identified 349 of such unique variants. 46 of them form 23 sets, such that mutations within each set are located within 100 bp of each other in the same clone, an event that is unlikely to happen by chance. Thus, we consider each such set as one “complex” mutation event^36,67^. The final set of detected mutations is listed in Table S3.

#### Comparison of LOH rates across strains

We define the LOH event rate as the number of LOH events per unit time (irrespectively of their length), and we define the LOH conversion rate as the probability per unit time that a position will be converted from heterozygous to homozygous state by an LOH event^46^. We estimate genome-wide and chromosome-arm-specific LOH event rates and LOH conversion rates as in Ref. ^46^.

To analyse finer-scale rate variation, we divide each chromosome into 50 kb non-overlapping windows as described in Ref. ^46^. (Note that Figures 3, <u>S6</u> and <u>S7</u> show results for overlapping 50 kb windows, which produces smoother profiles, but all statistical tests are done for non-overlapping windows.) We then calculate the per-basepair per generation event rate for each window for strain *s* as λ*_s_*= *k_s_*/*n_s_TL*, where *k_s_* is the number of LOH events detected in that window across all MA lines descendent from founder *s* corrected for undetected events (see above), *n_s_*is the number of these MA lines, *T* = 800 is number of generations, and *L* is the window size (50 kb for most windows). To estimate the 95% confidence intervals within each window and strain, we use the bootstrapping approach, i.e., we resample the number of events with replacement from their empirical distribution across clones.

To test whether the distributions of LOH events across the genome are statistically different between two founders, we calculate the difference in their cumulative distributions of LOH events across each chromosome using the Wasserstein statistic from the twosamples R package^68,69^. This statistic is calculated as the integral of the function |*E*_1_(*x*) – *E*_2_(*x*)|, where *E*_1_ and *E*_2_ are the empirical cumulative distribution functions for the two strains^70^. P-values are calculated from 5,000 iterations of a Monte Carlo simulation where we randomly reshuffle the founder labels of all LOH events.

To test for differences in genome-wide LOH conversion rates between two founders, we perform a permutation test on the mean fraction of the genome converted by reshuffling founder labels across LOH events. To test for differences in local LOH conversion rates, we perform permutation tests of the mean rate on each 50 kb window by reshuffling founder labels across clones. We then apply Benjamini-Hochberg correction with FDR ≤ 0.05 across windows for each founder comparison.

##### gRNA similarity

The probability that the Cas9:gRNA complex binds and cuts DNA at a given locus depends on the presence of a PAM site and on the 20-bp sequence immediately upstream of it^18,26,27^. To identify genomic regions where the cutting activity of the Cas9:gRNA complex might be elevated, we first index all genomic positions with either the canonical -NGG or the weakly interacting -NAG PAM sequences on both strands of the BY and RM reference genomes. Then, for each PAM position, we score the 20-bp region immediately upstream against our gRNA sequence using the empirically-informed scoring function `getMITScore` from the package crisprScore in R^18,71^.

To look for regions that may have elevated LOH rate in founder *s* = C or D compared to the W founder, we compute the normalized LOH event rate in window *i* as *r_si_* = (*R_si_* – *R_Wi_*)/(*R_si_* + *R_Wi_*), where *R_si_* and *R_Wi_* are the LOH event rates in window *i* in founder *s* and *W*, respectively. Windows are constructed as above, but with a slide interval of 50 kb such that windows are independent. We correlate this normalized rate with the maximum gRNA similarity score in each window (using the mean gRNA similarity score produces qualitatively similar results).

## Acknowledgements

We thank Shohreh Sikaroodi for help with experiments, Kryazhimskiy and Meyer lab for numerous helpful discussions. SK, JRM and OSA acknowledge support from DARPA (HR0011-17-2-0047), SK acknowledges support from NIH (R35GM153242). MSO was supported by the Pathways in Biological Sciences NIH T32 program (5T32GM133351-02). SEG was supported by the Tata Institute for Genetics and Society BS/MS Fellowship.

## Supplemental Figures

**Figure S1.**
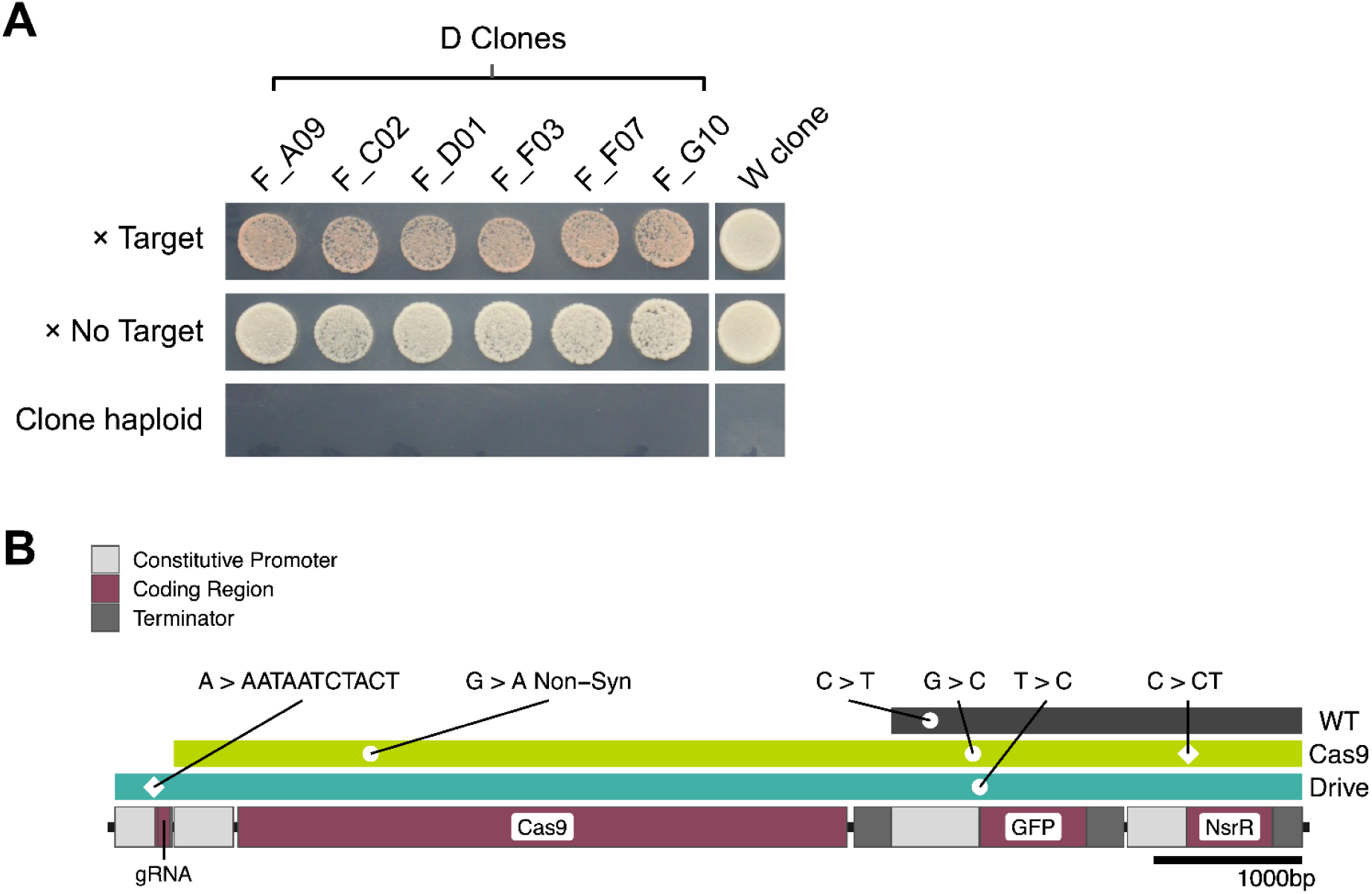
CCGD remains active throughout the MA experiment. **A.** All tested D end-point clones retain drive activity (see Section “<u>Testing of gene-drive activity</u>” in Materials and Methods). A haploid offspring of each tested D clone (columns) was mated either with a tester strain with the sequence targeted by the gRNA (Target) or with a tester strain without the target sequence (No target). Mated and unmated (“Clone haploid”) liquid cultures were spot plated on CSM + 5FOA agar, then replica plated on CSM + G418 agar for diploid selection and incubated for 48 hours. A successful disruption of the ADE2 gene by the CCGD results in red colony coloration^33^**. B.** Mutations found in the construct. The construct map shows the position of each component and the three colored bars above represent the portion of the construct integrated into each strain. Circles indicate point mutations and diamonds indicate indels.

**Figure S2.**
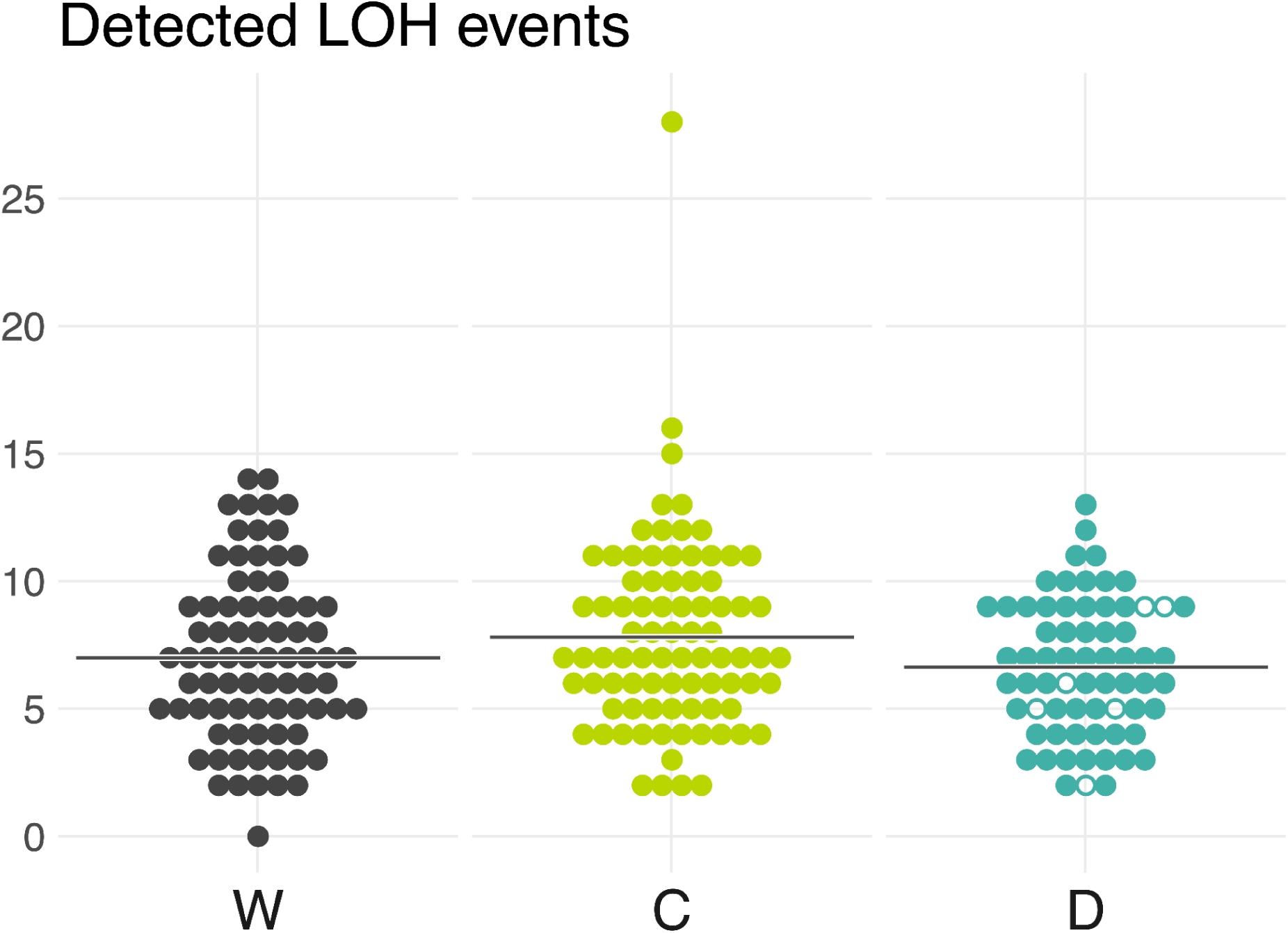
Distributions of detected LOH events across MA lines. Same as Figure 2B, but prior to undetected event corrections.

**Figure S3.**
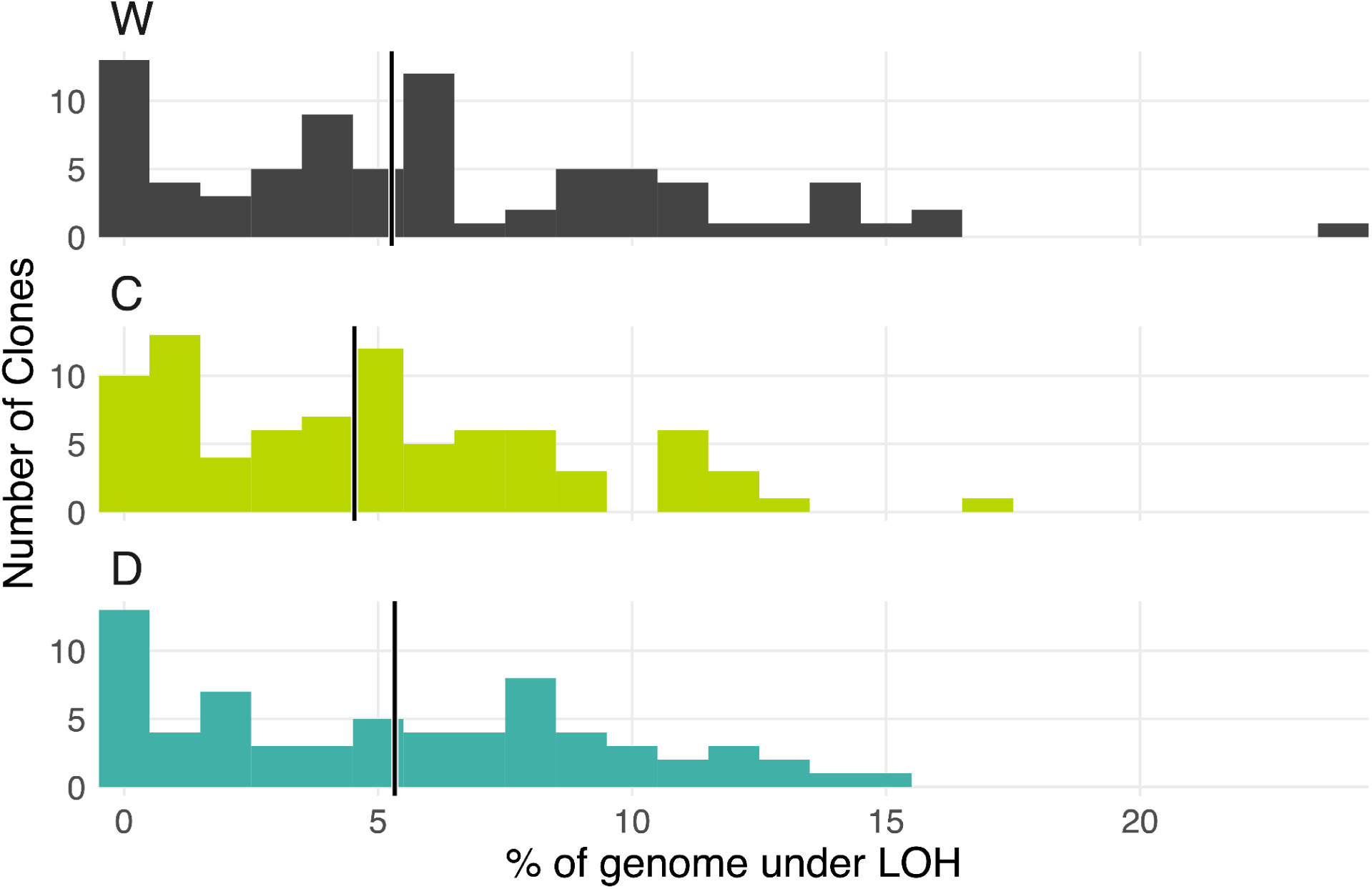
The rate of LOH conversion is not statistically elevated in CCGD element carrying strains.

**Figure S4.**
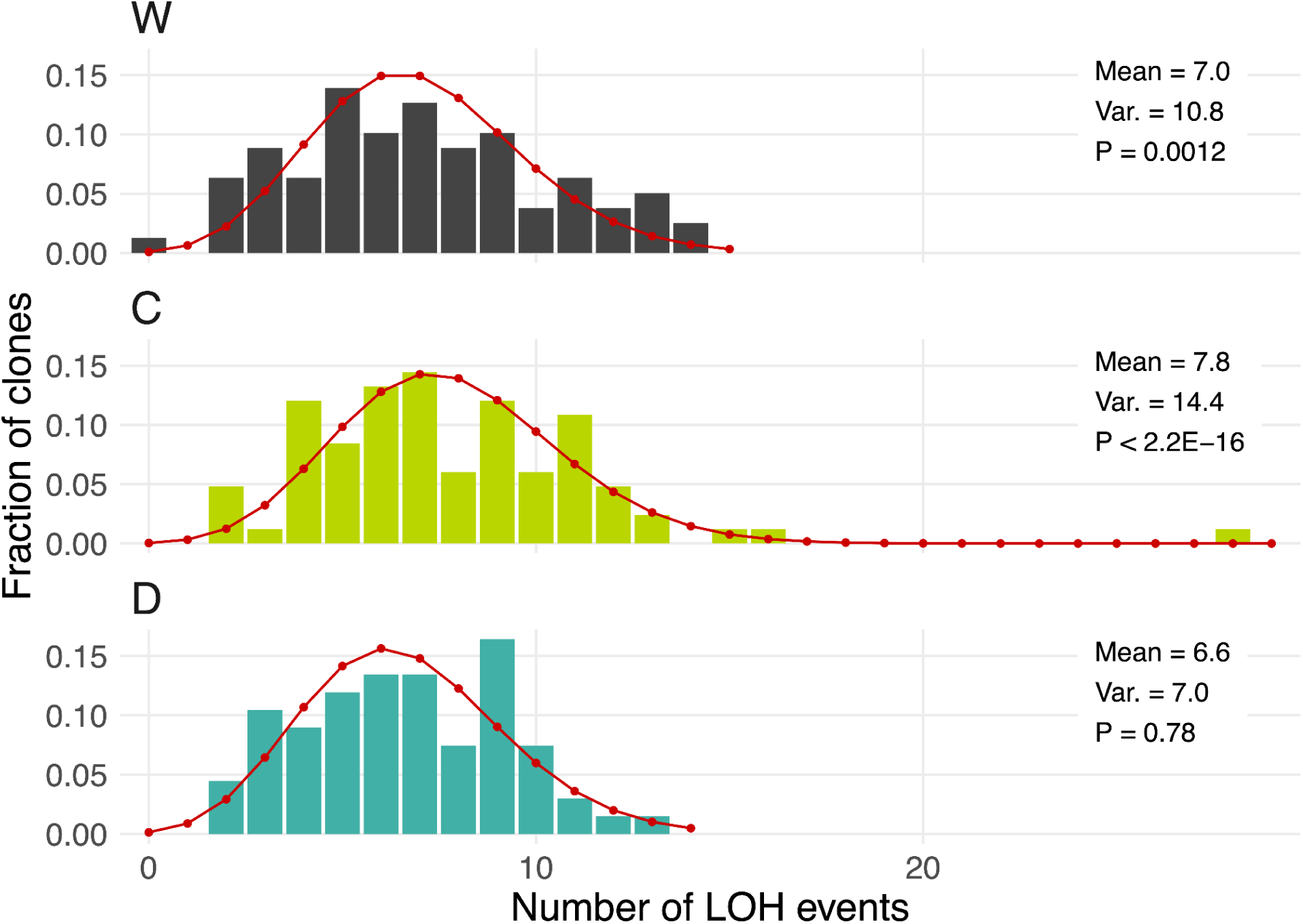
The distribution of LOH events across clones. The best-fit Poisson distribution is shown in red for each strain. *P*-value is calculated for the best-fit Poisson distribution based on the χ^2^ goodness-of-fit test.

**Figure S5.**
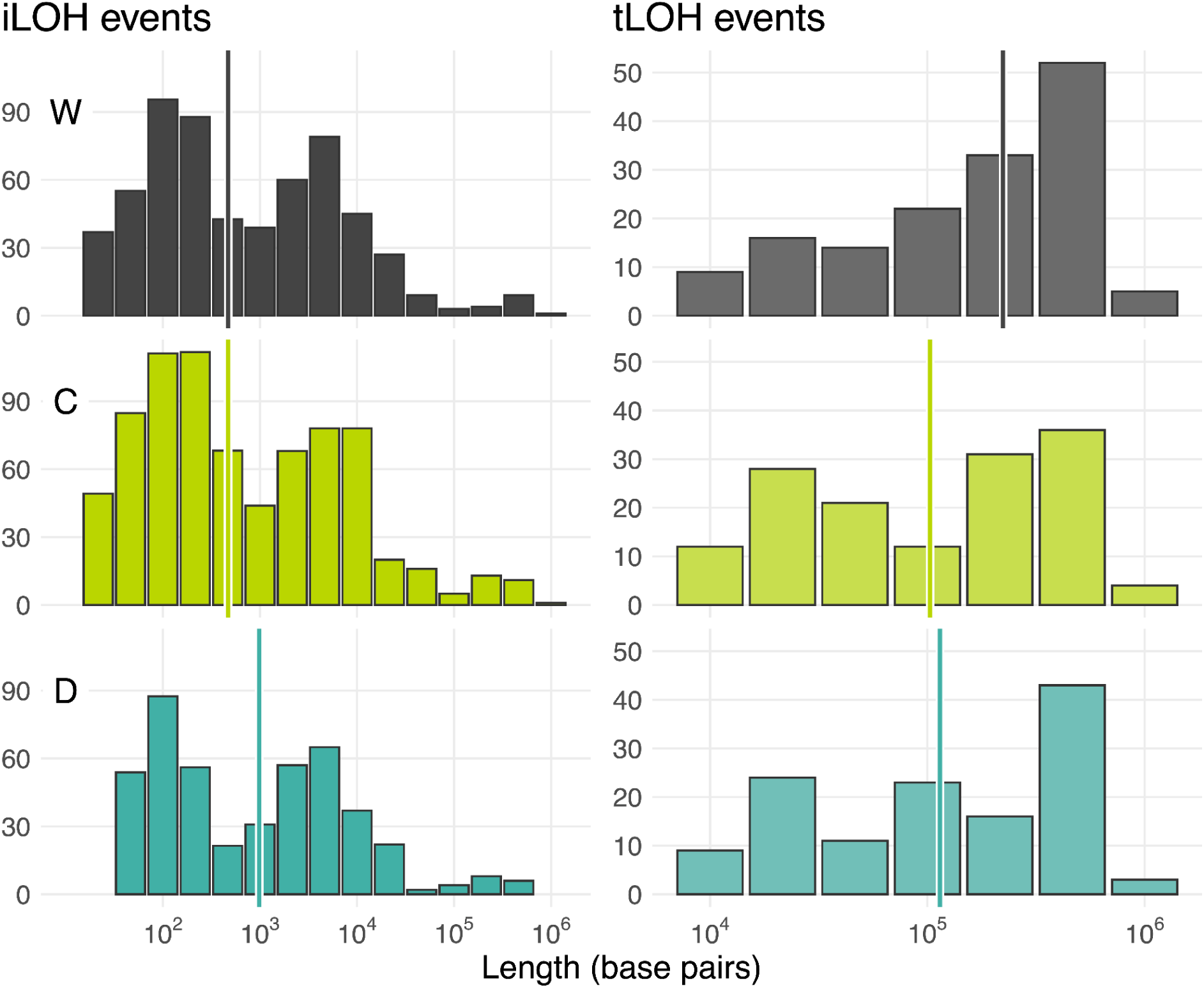
iLOH and tLOH length distributions across the three strains.

**Figure S6.**
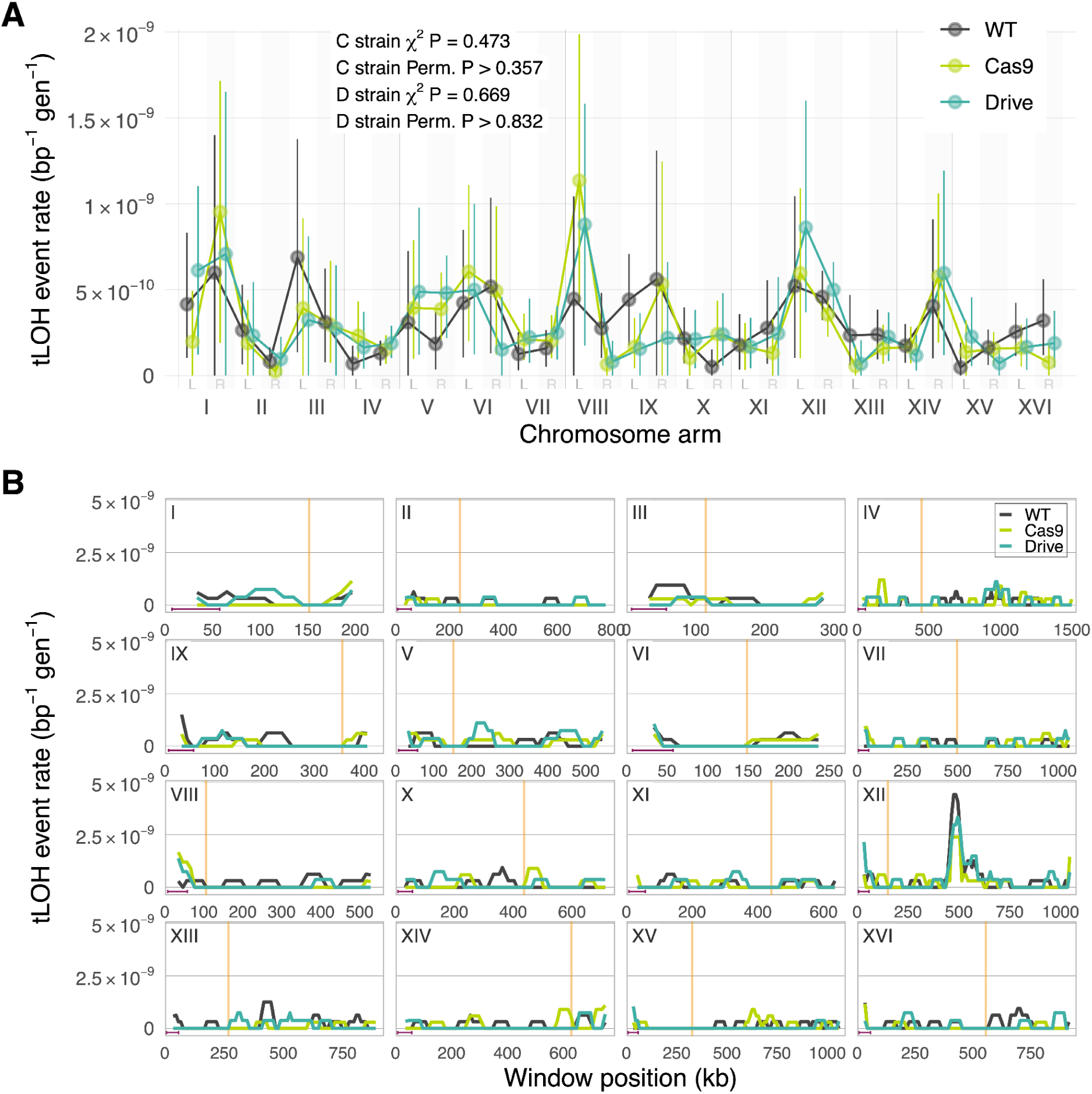
Distribution of tLOH event rates across the genome. Same as Figure 3, but for tLOH events.

**Figure S7.**
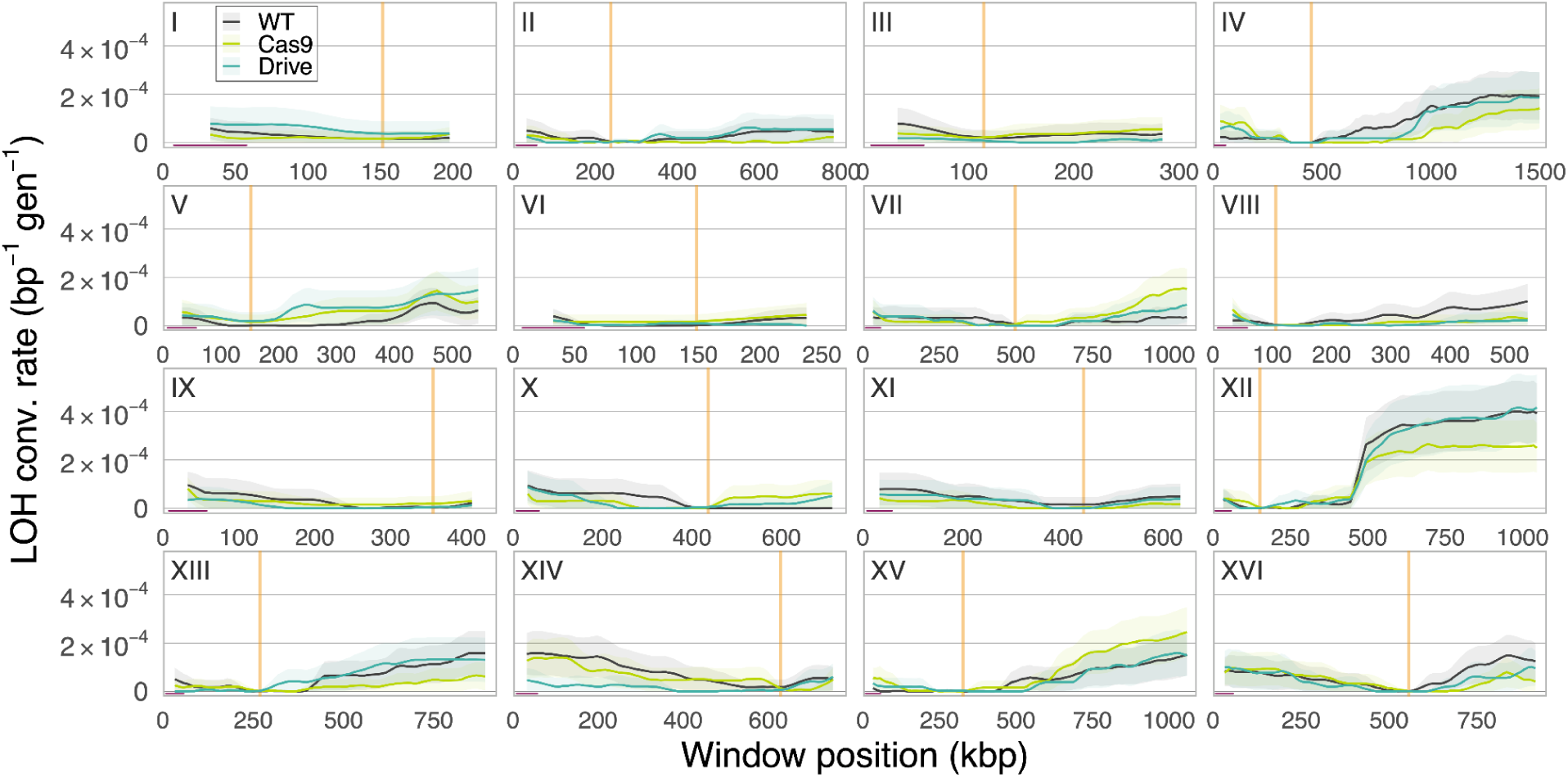
Distribution of LOH conversion rates across the genome. Each line shows the LOH conversion rate estimated within 50 kb sliding windows at 10 kb step sizes. Ribbons indicate 95% bootstrap confidence intervals. Maroon line segments are 50 kb scale bars, orange vertical lines are centromeres.

**Figure S8.**
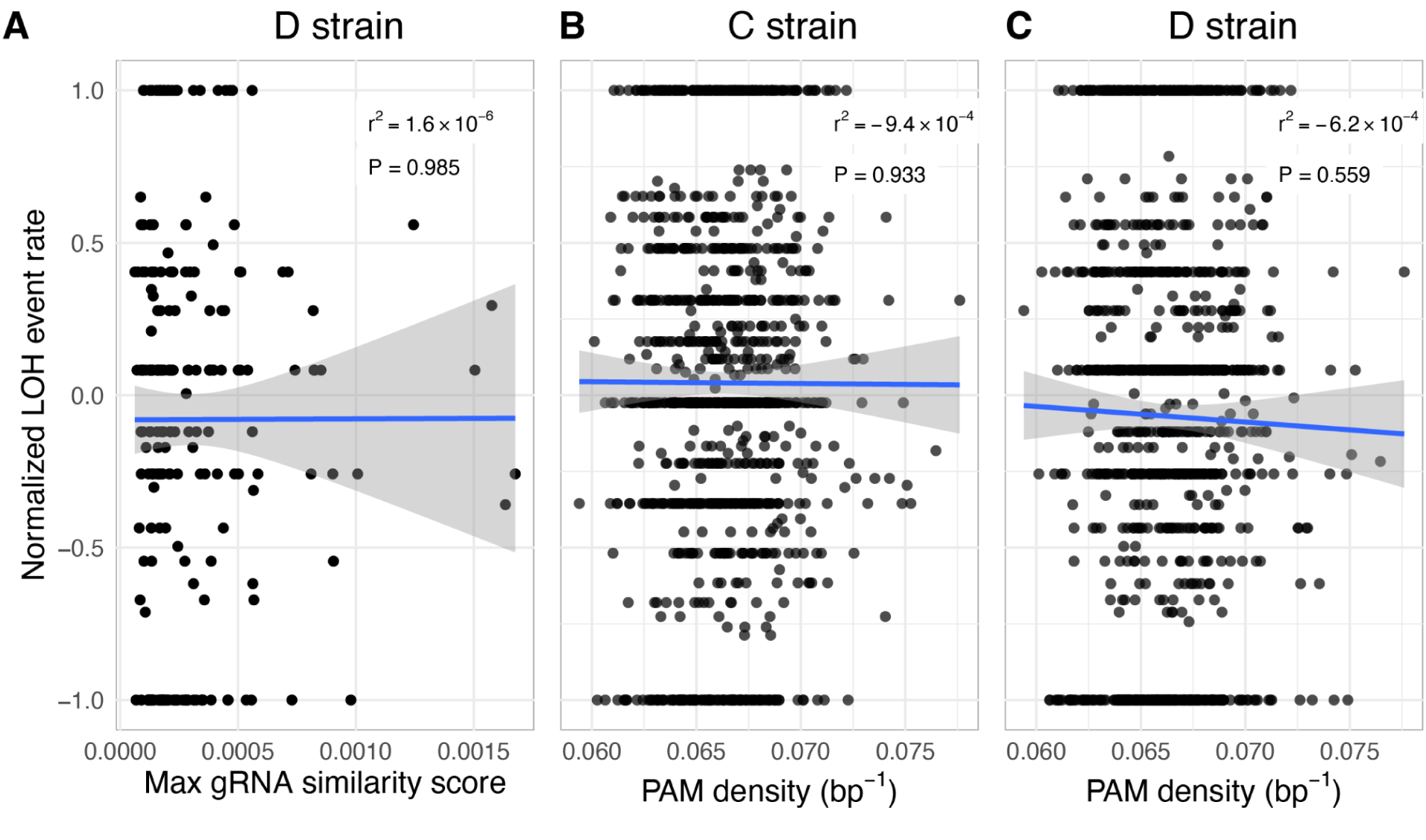
Correlation between the LOH event rate and sequence features across 50 kb genomic windows. Each point represents a 50 kb window in the genome. LOH event rates for C and D strains are normalized to the W rate in the same window as described in Section “<u>Comparison of local LOH rates</u> <u>across strains</u>” in the Materials and Methods. **A.** Normalized rate is plotted against the maximum similarity score between the PAM-adjacent genomic sequences and the gRNA found in a window. **B, C.** Normalized rate is plotted against the density of PAM sites in a window. For all panels, each point is a genomic window, blue lines are linear regressions, gray areas represent 95% confidence intervals.

